# Invariant scaling of the Species Abundance Distribution in observed and simulated marine plankton communities

**DOI:** 10.64898/2026.08.12.744266

**Authors:** Danling Ma, Enrico Ser-Giacomi, Yubin Raut, Stephanie Dutkiewicz, Oliver Jahn, Michael Follows, Gregory L. Britten

## Abstract

Marine plankton are functionally diverse and span over five orders of magnitude in diameter, with important consequences for marine biogeochemical cycles. Marine ecosystem simulations are beginning to resolve this diversity; however, major uncertainties persist regarding the structure and function of planktonic ecosystems. Here we diagnosed plankton Species Abundance Distributions (SADs) in large-scale surveys and in a global, mechanistic plankton community simulation. The fitted slopes of the SADs vary by less than 10% across latitude, season, and biome in both observations and the simulation. Fitting parametric SADs further reveals spatial structure in the shape and functional form of the SAD aligned with established biogeographic provinces. Together, these results demonstrate a largely invariant structure of marine plankton communities that persists despite strong environmental gradients and taxonomic turnover. These findings suggest that the emergent shape and scaling of plankton SADs reflect fundamental constraints on community assembly and provide a compact quantitative diagnostic for planktonic ecosystem structure.

## 1 Introduction

Marine plankton, including phytoplankton, zooplankton and bacterioplankton, form the foundation of ocean ecosystems, driving energy flow through food webs, sustaining higher trophic levels and mediating key biogeochemical cycles [1, 2]. Phytoplankton contribute approximately half of global net primary production [3, 4], while zooplankton regulate ecosystem functioning through grazing, nutrient recycling and carbon export [5–7]. Bacterioplankton remineralize organic matter, regenerating nutrients that sustain productivity and fuel higher trophic levels [8]. Together, these groups underpin the ocean’s biological pump and play a central role in climate regulation [9]. Yet, despite their importance, planktonic ecosystem structure remains incompletely resolved, with models often relying on highly simplified parameterizations of plankton biodiversity [10–12].

Spanning more than five orders of magnitude in diameter, from sub-micron bacteria to millimetre-scale zooplankton, the global plankton assemblage forms one of the most abundant and diverse living systems on Earth, with concentrations at times exceeding 10^6^ individuals per millilitre [13–16]. Large-scale observational and modelling efforts have begun to reveal how this extraordinary diversity is structured. The *Tara* Oceans expedition, for example, combined environmental and metagenomic data to map plankton interaction networks and carbon-export potential [17], while global niche models linked more than half a million observations to patterns of phytoplankton diversity [18]. Other studies have projected future biogeographic shifts under climate change [19], analysed global abundance distributions [20], and tested neutral theory against latitudinal gradients in diversity [21].

Species abundance distributions (SADs), which describe the relative abundance of species in a community, are a central macroecological descriptor of biodiversity and community structure. Operationally, an SAD is constructed by measuring the abundance of each taxon within a local community and summarizing the resulting abundances as a frequency distribution across abundance classes. A local community in this case corresponds to one surface microscopy sample, one station-matched simulation sample, or one model grid cell at a specified time or averaging period (see Methods). SADs commonly capture a characteristic feature of ecological systems: a small number of highly abundant taxa and a long tail of many rare species [22–24]. Because they retain information across the abundance spectrum, SADs provide a diagnostic of diversity, community evenness, and dominance, and have been widely 2 used to investigate emergent macroecological patterns and the processes underlying community assembly, including niche differentiation, stochasticity, and dispersal [24].

In marine plankton, Ser-Giacomi et al. [20] demonstrated emergent macroecological properties of protist (single-celled eukaryotes) communities in which SADs follow a nearly constant power-law exponent across ocean basins, suggesting a universal scaling structure within an individual taxon that persists despite environmental and taxonomic turnover. This apparent invariance echoes other macroecological patterns observed across marine and terrestrial systems; for example, the near-constant slope of the global biomass size spectrum linking bacteria to whales [16], the coupled scaling of abundance, body size and metabolism predicted by metabolic and allometric theory [25, 26], and the cross-system predictions of the Maximum Entropy Theory of Ecology [27]. At the same time, plankton communities differ from terrestrial systems because fluid transport continually introduces taxa into local assemblages, maintaining a pool of rare or non-dominant species. Previous interpretations of plankton SADs emphasized that transport-induced dispersal, together with demographic stochasticity, may contribute to the power-law structure of the non-dominant community in marine systems [20]. More broadly, SADs have been interpreted through alternative null models of community assembly. Neutral and dispersal-based frameworks explain abundance structure from ecological equivalence, stochastic birth and death, and immigration from a regional species pool [28–30]. Niche- or trait-based consumer–resource models contrast with neutral models in allowing taxa to differ in growth, grazing, mortality and resource acquisition, so that relative fitness differences shape community structure [31–33]. Thus, SAD invariance may reflect both local constraints on growth, mortality and trophic interactions, while neutral processes can stochastically shuffle and redistribute taxa across oceanic habitats.

Here we examine planktonic SADs in models and observations across broad oceanographic and taxonomic gradients. We test the generality of the Ser-Giacomi et al. results [20] by explicitly analyzing observations that span the broader taxonomy of marine plankton and compare patterns with a mechanistic plankton ecosystem model to (1) characterize scaling relationships in community planktonic observations across taxa, (2) compare observed scaling to those simulated from a mechanistic plankton model, and (3) use the model to predict SAD scaling across environmental gradients in a global context.

## 2 Results and Discussion

### 2.1 Consistent scaling of the species abundance distribution in observations and the simulation

Due to limited global observational coverage of plankton communities, we focus our empirical analysis on the Atlantic basin, which has been repeatedly sampled with consistent methodology across a wide latitudinal gradient, intersecting regions of high plankton biomass near equatorial upwelling zones and boundary currents and low biomass in subtropical gyres (Fig. 1 and Supplementary Table 1) [34]. These observations provide a representative baseline for characterizing empirical marine SADs and comparing against SADs derived from the simulation.

**Fig. 1.**
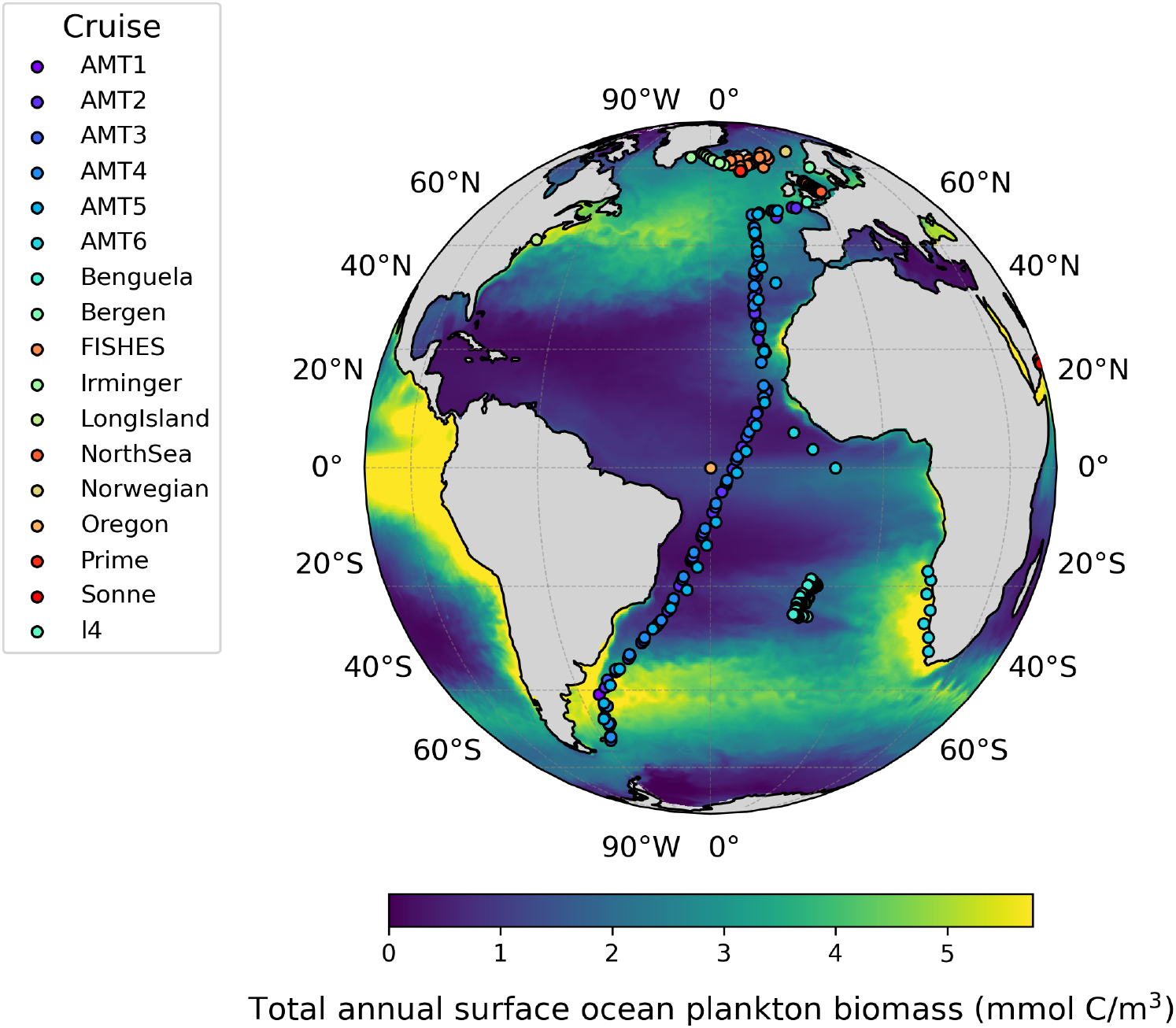
Study region map showing plankton sampling sites from the Marine Microplankton Diversity Database across the Atlantic Ocean [34]. Sampling points are overlaid on the simulated total annual surface ocean plankton biomass from the Darwin model (mmol C/m^3^). Sampling locations are color-coded by cruise ID.

We analyze a simulation from the MIT Darwin plankton ecosystem model [32, 33, 35–37] which is a trait-based plankton community model embedded within the MIT general circulation model (MITgcm) [38, 39]. The Darwin framework (hereafter simply ‘the simulation’) includes mechanistic plankton dynamics and the complex physical and biogeochemical ocean environment (see Methods). The explicit integration of ocean circulation, chemistry, and plankton community dynamics, including 35 phytoplankton and 16 zooplankton functional types, allows large-scale simulations in which plankton growth, mortality, and community assembly dynamically interact and feedback onto ocean biogeochemistry. By resolving functional diversity through a trait-based formulation, the model provides a testbed for understanding emergent macroecological community structure at the basin and global scales.

We first quantified the first-order SAD scaling in observations and the simulation by fitting linear regressions to the log–log relationship between frequency density and abundance (Figs. 2 and 3). Across all biomes, the simulation closely reproduces the observed abundance distributions, capturing both the linear scaling on log–log axes and the relatively narrow distribution of regression slopes across latitude, season, and biome (Supplementary Fig. 1). Globally, the mean slope of observed SADs (−1.21) was slightly more negative than that of the simulation (−0.95), expressed in units of Δ log_10_(frequency density) per Δ log_10_(abundance). After normalization by dividing by their respective mean slopes, fitted SAD slopes varied by only 7.2% in the observations and 4.8% in the simulation, on average, across latitude, season, and biome (Fig. 2). Within the global simulation, these normalized slopes remain nearly invariant across the global ocean and throughout the seasonal cycle (Supplementary Fig. 1). In the simulation, SAD slopes show little basin-scale structure when fitted to SADs constructed from both daily abundance and annual means. Normalized slopes remain close to the global mean slope across coastal, trade wind, and westerlies biomes, with little apparent spatial structure (Fig. 3b). Slope distributions are broadest in the polar biome in both observations and the simulation (*σ* = 0.089 and 0.065, respectively; Fig. 2f), where slopes also show the strongest temporal variability (Fig. 3c). We note that simulated SADs extend further into the high-abundance tail than the observations, reflecting the absence of the smallest cyanobacteria (e.g., *Prochlorococcus, Synechococcus*) from the microscopy dataset, where these taxa are not reliably identified [34]. We also note that both datasets exclude heterotrophic bacteria: the observational SADs are based on microscopy-resolved microplankton taxa larger than 10 *µ*m, while the simulated SADs include phytoplankton and zooplankton functional 6 types but not heterotrophic bacteria. The close agreement in regression slopes and overall SAD scaling indicates that the simulation captures the emergent and invariant scaling of planktonic SADs.

**Fig. 2.**
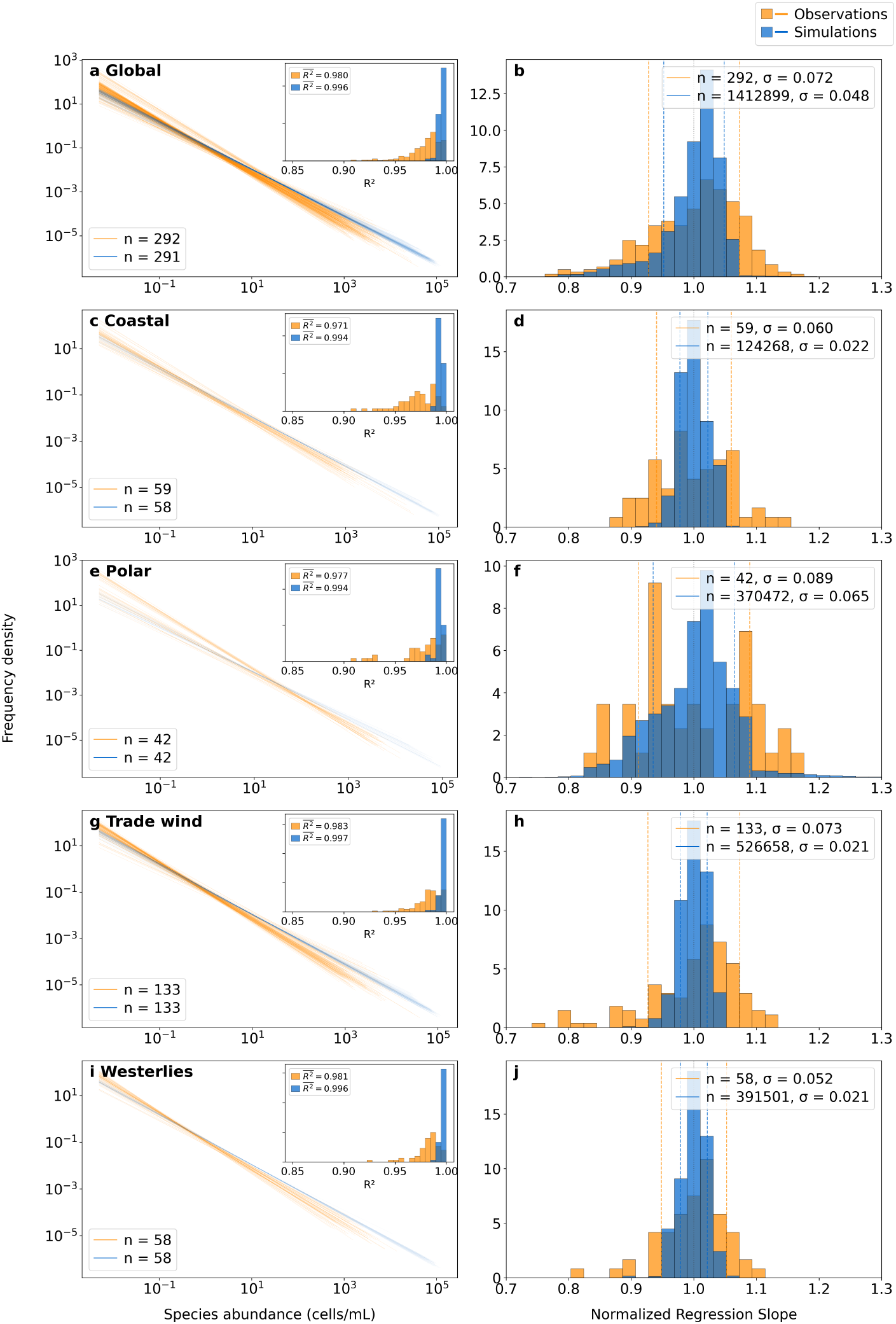
Regression analysis comparing observed and simulated species abundance distributions across the global ocean and major biomes. Observations are derived from surface microscopy counts across all sites in Figure 1. In the left panels (a, c, e, g, i), simulated SADs are station-matched to the observations by selecting the nearest model grid cell and corresponding day of year for each observational sample. These panels show log–log linear regressions of individual SADs for observations (orange) and station-matched simulated output (blue), with distributions of regression *R*^2^ values shown in the insets. In the right panels (b, d, f, h, j), observed slope distributions are compared with simulated slope distributions calculated from all model grid cells within the corresponding biome, so the simulated sample sizes are larger than in the station-matched comparison. Observed and simulated slopes were normalized separately: global slopes were divided by their respective global means, whereas slopes within each biome were divided by the corresponding observational or simulated biome mean. Dashed lines indicate 1 *± σ*.

**Fig. 3.**
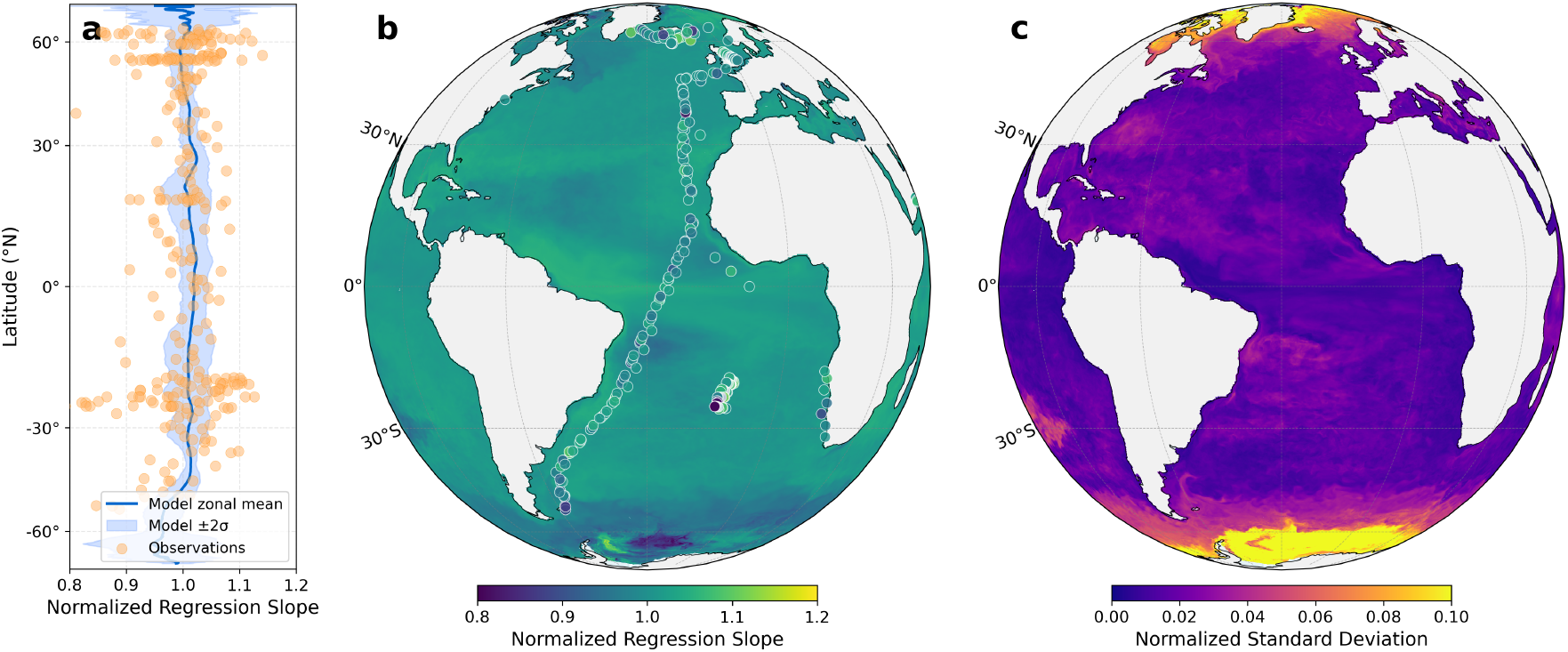
Spatial and temporal variability in normalized regression slopes of species abundance distributions. (a) Zonal average of normalized temporal-mean simulated slopes versus latitude, showing the simulation mean (blue) with *±* 2*σ* variation across longitude (shaded) and observations (orange circles). (b) Spatial distribution of temporal-mean simulated slopes, normalized by their global spatial mean. (c) Temporal standard deviation of slopes after normalization by the temporal-mean slope at each grid cell. Slopes were calculated at each time step from log–log regressions of abundance-frequency relationships. For panels a and b, the temporal-mean simulated slope at each grid cell was normalized by the mean of these slopes across all valid simulated ocean grid cells. For panel c, time-specific slopes were normalized by the temporal-mean slope of the corresponding grid cell before their temporal standard deviation was calculated. Observational points were normalized by the corresponding cruise mean slope.

Observations and the simulation indicate that the invariance of SAD scaling, first identified among non-dominant protist taxa [20], persists across communities spanning many taxa, multiple trophic levels, size classes, and environmental regimes. By combining observations and a global simulation, our analysis generalizes this earlier observation based study of plankton SADs, suggesting that invariant SAD scaling is a pervasive feature of marine planktonic communities, reflecting a broad constraint on community structure that is insensitive to local environmental variability and community composition.

To better understand how SAD scaling emerges from mechanistic assumptions, we conducted idealized simulations using a simpler non-spatial size-structured plankton model from Armstrong (1994) [31]. We simulated a size-structured plankton community including 50 phytoplankton and 50 zooplankton while systematically varying the allometric distribution of growth rates in the community. We found that SAD scaling depended strongly on the assumed allometric parameters: varying the allometric exponent for maximum phytoplankton growth rate, 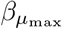, shifted the fitted SAD slopes across a wide range, including changes in slope sign (Supplementary Fig. 2).

This sensitivity suggests that invariant SAD scaling is not a generic outcome of size-structed consumer–resource dynamics, but instead requires additional constraints on the community dynamics, including growth rate distributions, but perhaps also spatial processes including dispersal and resource transport. The contrast with the Darwin simulation is therefore informative. Unlike the idealized simulations, the Darwin configuration embeds fixed allometric trait relationships, with scaling exponents drawn from published empirical studies, within a three dimensional circulation and biogeochemical framework whose physiological parameterizations have been developed to reproduce observed large scale patterns in nutrients, chlorophyll, plankton biogeography, and biogeochemical fluxes [32, 33, 37, 40, 41], leading to emergent SAD invariance. Importantly, the simulation was not tuned to reproduce SAD slopes or fitted SAD forms. We suggest that future theoretical work should investigate the conditions under which idealized ecosystem models reproduce invariant SAD scaling, including the relative roles of trophic complexity, dispersal, trait diversity, and parameter structure. Such analyses are beyond the scope of the work presented here.

### 2.2 Biogeographical patterns

The log–log regressions show that SADs share a broadly invariant first-order scaling structure; however some scatter remains unexplained by the log–log linear fits. We therefore asked whether variation not captured by the slope was organized biogeographically by fitting and comparing parametric SAD forms across biomes. We compared multiple candidate probability density functions (PDFs) that represent alternative hypotheses about community assembly processes, including lognormal distributions linked to the cumulative effect of many multiplicative processes or niche partitioning, and power-law or truncated power-law forms associated with scale-invariant dynamics, stochastic demographic processes, and dispersal-driven assembly [20, 22, 24, 42]. By comparing these candidate PDFs, we do not assume a single “true” generating process, but rather assess if the shape of the SAD varies systematically across oceanographic gradients, since different ecological processes can generate similar SAD functional forms [20, 24]. We fitted five candidate PDFs to simulated SADs across the global ocean grid, following Longhurst biogeographic provinces [3]. We compared lognormal [22], power law [42], truncated power law [20], Ewens sampling formula [43], and the meta zero sum multinomial [30], using the Akaike information criterion (AIC) for PDF selection [44] (see Methods).

Mapping best fit PDFs across the global ocean grid reveals coherent large scale biogeographic structure within the simulation (Fig. 4; Supplementary Fig. 3). Lognormal SADs dominate the coastal and westerlies biomes, accounting for 74.7% and 72.1% of fitted grid cells, respectively. The polar biome is instead most frequently characterized by truncated power laws, which account for 65.8% of fitted grid cells. The trade wind biome is intermediate, with similar support for lognormal and power law PDFs, which account for 51.3% and 47.2% of fitted grid cells, respectively. Neither the Ewens sampling formula nor the metaZSM emerges as conclusive best fits.

**Fig. 4.**
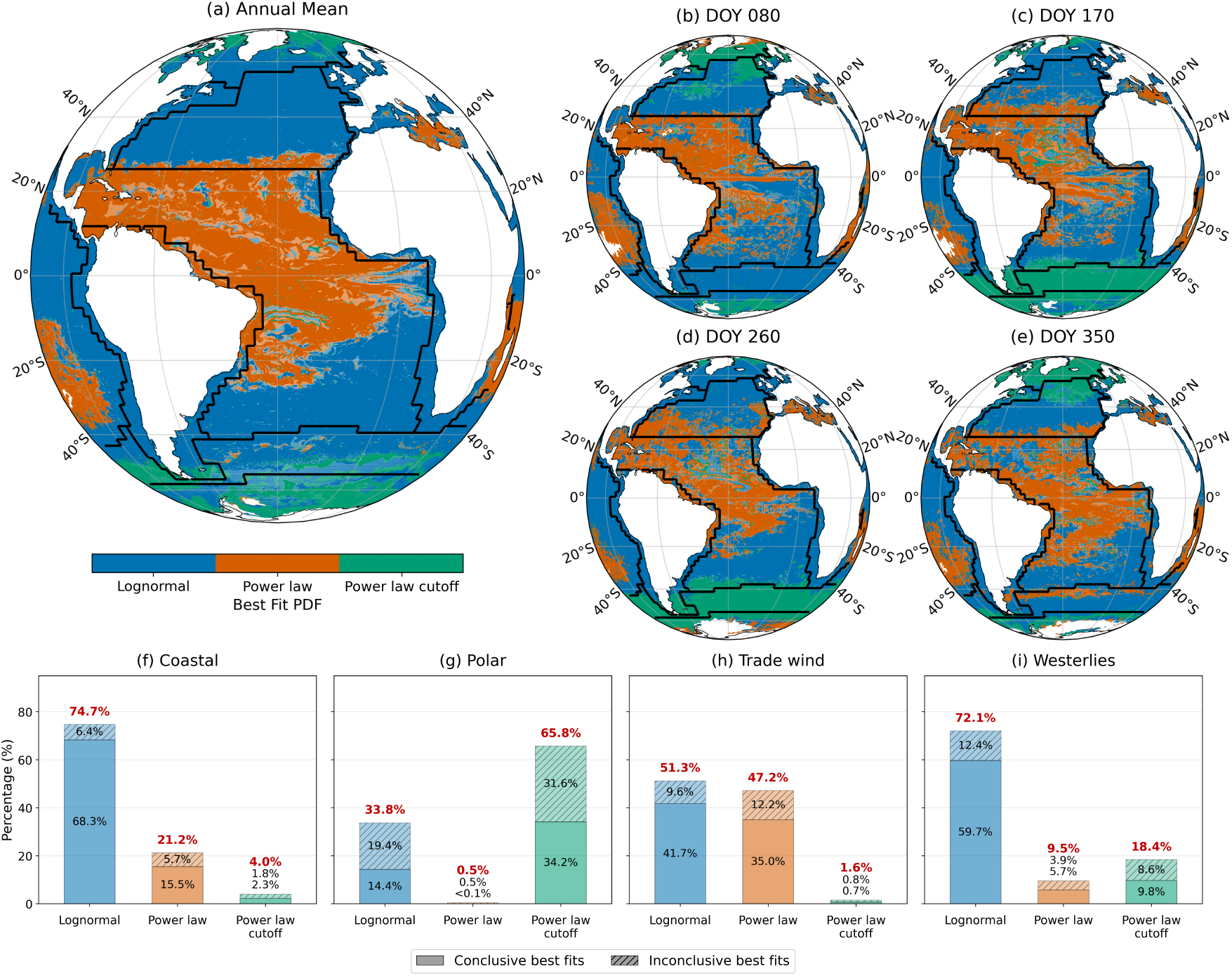
Spatial distribution of the best fitting probability density function (PDF) for simulated species abundance distributions and biome-scale best fitting PDF percentages across all model grid cells. (a) Annual mean classification at each grid cell. (b–e) Seasonal snapshots showing the best fitting PDF on day of year (DOY) 80, 170, 260, and 350, respectively. Colors indicate the PDF type with lowest AIC: blue for lognormal, orange for power law, and green for power law with cutoff. Lighter shades in the maps denote inconclusive cases where ΔAIC *<* 2 between the two best supported PDFs. Black contours show biome boundaries. (f–i) Percentage of all Darwin model grid cells best fit by each PDF type within the Coastal, Polar, Trade wind, and Westerlies biomes, respectively. Conclusive fits (ΔAIC ≥ 2) are shown as solid bar segments, while inconclusive fits (ΔAIC *<* 2) are hatched and stacked above the corresponding conclusive segments. Black percentage labels indicate the individual conclusive and inconclusive contributions, and bold red labels indicate the total percentage for each PDF type.

The polar dominance of truncated power laws suggests that the SAD retains a long tail of rare or low abundance taxa, but contains fewer extremely abundant taxa than expected under an unbounded power law. One possible explanation is that strong seasonality, mixed layer deepening, sea ice retreat, bloom initiation, and episodic transport disturb local communities, limiting the duration over which any one taxon can remain strongly dominant, while dispersal and mixing may maintain a pool of rare or non dominant taxa. This interpretation is broadly consistent with previous work showing that phytoplankton communities reflect both niche and neutral processes, and that inferred immigration rates increase poleward, likely because of stronger mixing and stratification cycles [21]. The dominance of lognormal SADs in the temperate westerlies biome may reflect communities shaped by multiple interacting processes, including seasonal succession, resource partitioning, and variable growth and loss rates, which can produce a broad distribution of taxa at low to intermediate abundances. Similarly, lognormal fits in coastal regions may reflect the combined influence of nutrient supply, mixing, grazing, and habitat heterogeneity, which can generate multiplicative variation in taxon abundances across local communities. By contrast, SADs in the relatively stable, weakly seasonal trade wind biome are more often described by power laws, consistent with persistent abundance hierarchies in which a small number of taxa dominate while many others remain rare. These spatial patterns demonstrate that the simulation captures not only the global prevalence of lognormal and power law type SADs, but also their systematic biogeographic variation in SAD form across biomes. We repeated the same PDF fitting analysis for the observation matched samples and their corresponding station matched simulations (Supplementary Fig. 4). These comparisons show broad agreement where observations and the simulation overlap, with fitted SADs again concentrated within the lognormal and power law families. However, the observational samples are unevenly distributed within several biomes, so biome-aggregated observational fits are biased toward sampling locations rather than representing the dominant structure of the biome as a whole. This is especially relevant for the polar biome, where observations sample a small fraction of the polar domain. Where the simulation and observations overlap, best fitting SAD shapes are broadly consistent, indicating a coherent relationship between SAD shape and oceanographic regime (Supplementary Fig. 4).

It is important to note that observed and simulated SADs are defined at different levels of taxonomic resolution, where observations comprise microscopy-resolved taxa at (or near) the species level, while the simulation represents size-classes within plankton functional types. This mismatch is an unavoidable limitation as it is not computationally feasible to develop models at the species level, even if there were enough laboratory studies to assign the appropriate parameters. However, SAD agreement between observations and the simulation also provides evidence for a more basic set of structural constraints acting on planktonic ecosystems independent of the level of aggregation. Aggregation may help explain the slight offset in mean slopes between observations and the simulation despite similar invariance in slope variability. We again suggest further theoretical work to understand how SAD structure may vary across levels of taxonomic resolution.

## 3 Conclusion

We evaluated SADs in observations and a mechanistic simulation, finding that SADs of marine plankton communities exhibit near invariant first-order scaling and coherent biogeographical structure in parametric form. Across the observational dataset and the simulation, SAD slopes vary by less than 10%, on average, across latitude, season, and biome, extending earlier findings of invariant scaling in marine protists [20]. At the same time, variation unexplained by linear log–log scaling follows consistent biogeographical structure with lognormal forms most common in coastal and westerlies regions, truncated power laws most common in polar regions, and mixed support for lognormal and power-law forms in the trade wind biome. Thus, plankton SADs combine a broadly invariant first-order scaling relationship with biome-dependent variation in underlying distributional shape. These results pave the way for future theoretical development to understand how SAD scaling and shape emerge from the combined influence of environmental variability, demographic processes, species interactions, trait tradeoffs, dispersal, and levels of taxonomic aggregation [24, 27, 45–48]. Furthermore, we suggest that observed departures from SAD invariance, such as increased variability in scaling or shifts in SAD biogeography, may indicate changes in the drivers of planktonic ecosystem structure related to climate change, species invasions, or other natural and anthropogenic perturbations.

## 4 Methods

### 4.1 Observations

Cell count data for microplankton were obtained from a global database encompassing 788 stations sampled between 1992 and 2002 [34]. Counts were made using the Utermöhl inverted microscopy method [49], which enumerates cells larger than 10 µm but excludes picoplankton such as *Prochlorococcus* and *Synechococcus*. Thus, observed SADs represent microscopy-resolved marine microplankton, and larger nanoplankton, including diatoms, coccolithophores, dinoflagellates, flagellates, heterotrophic dinoflagellates, heterotrophic flagellates, and ciliates, where identifiable. The samples do not quantitatively measure picoplankton or heterotrophic bacteria. All identifications were carried out by a single expert taxonomist to ensure consistency. We restricted analyses to surface samples, where the minimum abundance threshold is 0.005 cells mL^−1^. We focused on the analysis of microscopy data instead of amplicon and genomic-based estimates due to known biases arising from differences in PCR efficiency [50], primer bias [51], and genomic copy number variation [52], which is particularly challenging when comparing abundance across broad taxonomic groups.

### 4.2 Simulation

The simulation analyzed in this study was generated with the MIT Darwin ecosystem model using the CS510 configuration, embedded within the MIT general circulation model (MITgcm) [38, 39, 53]. The CS510 configuration resolves ocean circulation at a nominal horizontal resolution of approximately 18 km with 50 vertical levels [40]. At this resolution, the model is eddy-permitting, allowing a coarse representation of mesoscale variability that influences nutrient transport and plankton community structure.

The Darwin model includes biogeochemical cycling of carbon, nitrogen, phosphorus, silica, iron and oxygen, as well as a dynamic plankton community. Plankton mediate the biogeochemical cycling of nutrients and are represented as a trophic network with distinct functional groups and allometric size-based feeding interactions, growth rates, and nutrient affinities [32, 33, 37, 54, 55]. Key physiological properties such as maximum growth rate, nutrient affinity, and light utilization vary systematically as a function of size within functional types. The simulation resolves 35 phytoplankton and 16 zooplankton functional types within a global circulation framework [40, 41].

We analyzed annual mean and day-of-year surface abundances from the year 2010 to characterize both mean spatial patterns and temporal variability in SAD structure. Simulated biomass was converted to cell counts using type-specific estimates of biovolume and carbon to volume ratios consistent with the observational dataset [56]. A plankton type is considered present in a given grid cell if the simulated biomass is above the minimum detection limit of the observations.

### 4.3 Construction of species abundance distributions

For each observed sample or simulated grid cell, we constructed a SAD from the abundance vector across taxa or simulated plankton functional types within that local community. Observed SADs were constructed from surface microscopy samples, with taxon abundances expressed in cells mL^−1^. Station-matched simulated SADs were constructed from the phytoplankton and zooplankton functional type abundances at the model grid cell and day of year corresponding to each observed sample. Simulated SADs were constructed individually for each surface model grid cell. For annual mean analyses, we first averaged the surface abundance of each simulated plankton functional type over the year and then constructed an SAD from the resulting annual mean abundances.

For all observed and simulated SADs, abundances below *x*_min_ = 0.005 cells mL^−1^ were excluded before regression analysis and frequency distribution fitting, which corresponds to the minimum detection limit of the microscopy measurements. The 11 resulting abundances were used directly for maximum likelihood fitting of candidate probability distributions and were converted to a log-spaced frequency-density histogram for regression slope estimation.

### 4.4 Regression slope estimation and normalization

For each observed sample or simulated grid cell, we quantified SAD scaling by first representing the abundances as a frequency-density histogram on a logarithmic abundance axis. The number of bins was chosen after evaluating standard data-driven histogram rules on log-transformed abundances, including Scott’s and Freedman–Diaconis rules [57, 58]. We used the median recommended bin number across these rules, corresponding to five logarithmically spaced initial abundance bins, for all regression analyses. This fixed scheme provides a consistent basis for comparing slopes across observations and the simulation while limiting empty bins in communities with relatively few retained taxa or functional types.

For each SAD, the bin edges spanned from the abundance cutoff *x*_min_ = 0.005 cells mL^−1^ to the maximum abundance in that sample or grid cell. Frequency densities were estimated using the histogram counts divided by total counts and bin widths. Empty bins were removed before fitting. Regressions with fewer than three non-empty bins were excluded. We fitted a linear regression to the log–log relationship between abundance-bin frequency density and abundance-bin center,

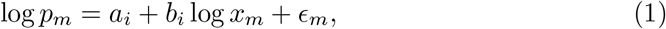

where *b*_*i*_ is the SAD regression slope for sample or grid cell *i, x*_*m*_ is the abundance-bin center, and *p*_*m*_ is the estimated frequency density. The regression *R*^2^ was recorded as the coefficient of determination of this log–log fit.

Observed samples and simulated grid cells were assigned to the four major ocean biomes (Polar, Westerlies, Trade Winds, and Coastal) using the Longhurst Provinces spatial dataset (Flanders Marine Institute, 2009; https://www.marineregions.org/). To compare slope variability across datasets, regions, and times, we expressed slopes as dimensionless ratios relative to a reference mean slope. For an observed or simulated slope *b*_*i*_ representing either the global ocean or a particular biome, the normalized slope was

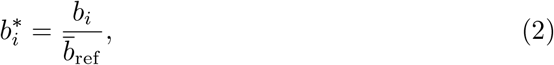

where 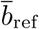 is the mean calculated from the observations or the simulation over the same spatial region as *b*_*i*_. For biome-resolved slope distributions, observed and simulated slopes were normalized separately: global slopes were normalized by the global observational or simulated mean, whereas slopes within each biome were normalized by that biome’s observational or simulated mean, respectively.

For spatial maps of simulated slopes, slopes were normalized by the global simulated mean slope so that map values show deviations from the global simulated SAD scaling. Observational points shown on the spatial summaries were normalized by their corresponding cruise mean.

For temporal variability maps, we first normalized the simulated slope at each time step, *b*_*t,i*_, by the temporal-mean slope at the same grid cell, 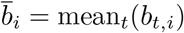. We then calculated the standard deviation of these normalized slopes through time:

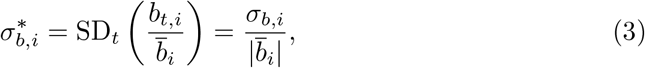

where *σ*_*b,i*_ = SD_*t*_(*b*_*t,i*_) is the temporal standard deviation of the unnormalized slopes at grid cell *i*. Thus, 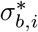 expresses temporal slope variability relative to the temporal-mean slope magnitude at each grid cell.

Because SAD regression slopes are negative, normalized values 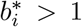 indicate slopes that are more negative, or steeper, than the reference mean, whereas 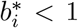 indicates slopes that are less negative, or flatter, than the reference mean. Standard deviations of normalized slopes are dimensionless; 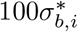 can be interpreted as the relative spread of slopes as a percentage of the reference mean slope. These normalized standard deviations describe variability in SAD scaling, whereas regression *R*^2^ describes the goodness of fit of each log–log linear SAD regression.

### 4.5 Fitting species abundance distributions

We evaluated SADs by fitting five candidate PDFs: the lognormal, the power law, the power law with exponential cutoff, the Ewens sampling formula (ESF), and the meta–zero–sum multinomial (metaZSM). The first three distributions were fitted directly to observed and simulated abundances by maximum likelihood estimation (MLE). Fits were applied to the abundance vectors described above. MLE finds the parameter set 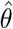 that maximizes the likelihood of abundances 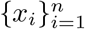,

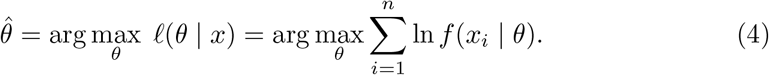

Closed–form estimates were used where available (e.g. power law exponent); otherwise we used gradient–based optimization. Fits were retained only for samples and grid cells with at least ten taxa after applying the abundance cutoff threshold, and only when the fitting procedure returned finite, admissible parameter estimates: 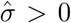 for the lognormal, 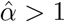 for the power law, finite 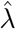 and 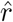 for the power law with exponential cutoff, and positive 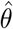 for the ESF and metaZSM fits.

The functional forms are as follows. The lognormal PDF [22] is given as

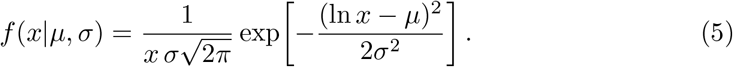

The power law PDF [42] is

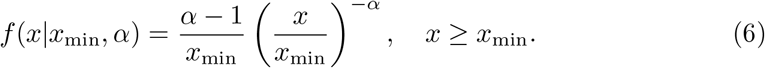

The power law PDF with an exponential cutoff [20] is

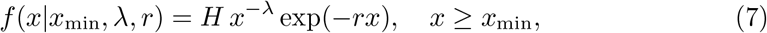

where the normalization constant was

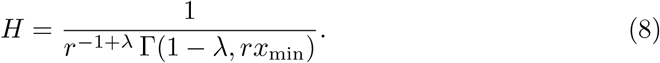

The Ewens sampling formula (ESF) [28, 43, 59, 60] gives the probability of an abundance vector (*x*_1_, …, *x*_*S*_) as

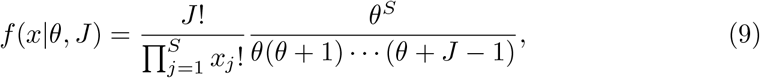

where 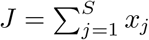 is the number of individuals, *S* is the number of species, and *θ* is the biodiversity parameter.

For the meta zero sum multinomial (metaZSM) [30], we fitted both the exact likelihood and the large *J* approximation. The two forms were indistinguishable across our datasets, consistent with the large number of plankton individuals. The approximate form is

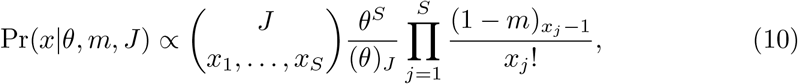

where *m* is the immigration rate, (*θ*)_*J*_ is the Pochhammer rising factorial, and (*a*)_*n*_ is the falling factorial. This form generalizes the ZSM by coupling the local community to a metacommunity via immigration.

### 4.6 Selection of SAD PDFs

The fit of candidate SAD PDFs was evaluated using the Akaike Information Criterion (AIC), which penalizes complexity while rewarding goodness of fit [61]:

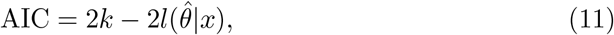

where *k* is the number of parameters in the fitted PDF and 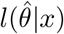 is the log likelihood evaluated at the maximum likelihood estimate.

Among the candidate PDFs (lognormal, power law, truncated power law, Ewens sampling formula, and meta–zero–sum multinomial), the PDF with the lowest AIC was selected as the best-supported description of the SAD. If the difference between the lowest and second-lowest AIC values was less than two, the result was considered statistically inconclusive.

## Supporting information

Supplementary Information for: Invariant scaling of the Species Abundance Distribution in observed and simulated marine plankton communities

## Supplementary information

Supplementary information is available as a separate file.

## Data availability

The Marine Microplankton Diversity Database analysed in this study is publicly available through Figshare [62]. The Longhurst Provinces spatial dataset is available from Marine Regions (https://www.marineregions.org/). The compact analysis ready data required to reproduce all main and supplementary figures are available in the GitHub repository (https://github.com/danlingmaa/marine-plankton-sad). The full time-resolved MIT Darwin surface biomass output analyzed in this study will be archived in a public repository and assigned a Digital Object Identifier upon acceptance of the manuscript.

## Code availability

All code used to preprocess the data, perform the statistical analyses, run the compute-intensive grid-cell workflows, and generate the main and supplementary figures is available in the GitHub repository (https://github.com/danlingmaa/marine-plankton-sad).

## Acknowledgements

DM and GLB were supported by Simons Foundation grant LS-CBIOMES-01195558. YR, SD, OJ, and MF were supported by Simons Foundation grant 549931FY22.

## References

[1] Lalli, C. & Parsons, T. R. Biological oceanography: an introduction 2nd edn (Butterworth-Heinemann, Oxford, UK, 1997).

[2] Falkowski, P. G., Barber, R. T. & Smetacek, V. Biogeochemical controls and feedbacks on ocean primary production. Science 281, 200–206 (1998).

[3] Longhurst, A., Sathyendranath, S., Platt, T. & Caverhill, C. An estimate of global primary production in the ocean from satellite radiometer data. Journal of Plankton Research 17, 1245–1271 (1995).

[4] Field, C. B., Behrenfeld, M. J., Randerson, J. T. & Falkowski, P. Primary production of the biosphere: integrating terrestrial and oceanic components. Science 281, 237–240 (1998).

[5] Calbet, A. & Landry, M. R. Phytoplankton growth, microzooplankton grazing, and carbon cycling in marine systems. Limnology and Oceanography 49, 51–57 (2004).

[6] Schmoker, C., Hernández-León, S. & Calbet, A. Microzooplankton grazing in the oceans: impacts, data variability, knowledge gaps and future directions. Journal of Plankton Research 35, 691–706 (2013).

[7] Steinberg, D. K. & Landry, M. R. Zooplankton and the ocean carbon cycle. Annual Review of Marine Science 9, 413–444 (2017).

[8] Azam, F. & Ammerman, J. W. in Cycling of organic matter by bacterioplankton in pelagic marine ecosystems: microenvironmental considerations (ed. Fasham, M. J. R.) Flows of energy and materials in marine ecosystems: theory and practice 345–360 (Plenum Press, New York, NY, 1984).

[9] Sarmiento, J. L. & Gruber, N. Ocean biogeochemical dynamics (Princeton University Press, Princeton, NJ, 2006).

[10] Franks, P. J. Planktonic ecosystem models: perplexing parameterizations and a failure to fail. Journal of Plankton Research 31, 1299–1306 (2009).

[11] Eyring, V. et al. Overview of the Coupled Model Intercomparison Project Phase 6 (CMIP6) experimental design and organization. Geoscientific Model Development 9, 1937–1958 (2016).

[12] Henson, S. A., Cael, B., Allen, S. R. & Dutkiewicz, S. Future phytoplankton diversity in a changing climate. Nature Communications 12, 5372 (2021).

[13] Massana, R., Balagué, V., Guillou, L. & Pedrós-Alió, C. Picoeukaryotic diversity in an oligotrophic coastal site studied by molecular and culturing approaches. FEMS Microbiology Ecology 50, 231–243 (2004).

[14] De Vargas, C. et al. Eukaryotic plankton diversity in the sunlit ocean. Science 348, 1261605 (2015).

[15] Lombard, F. et al. Globally consistent quantitative observations of planktonic ecosystems. Frontiers in Marine Science 6, 196 (2019).

[16] Hatton, I. A., Heneghan, R. F., Bar-On, Y. M. & Galbraith, E. D. The global ocean size spectrum from bacteria to whales. Science Advances 7, eabh3732 (2021).

[17] Guidi, L. et al. Plankton networks driving carbon export in the oligotrophic ocean. Nature 532, 465–470 (2016).

[18] Righetti, D., Vogt, M., Gruber, N., Psomas, A. & Zimmermann, N. E. Global pattern of phytoplankton diversity driven by temperature and environmental variability. Science Advances 5, eaau6253 (2019).

[19] Benedetti, F. et al. Major restructuring of marine plankton assemblages under global warming. Nature Communications 12, 5226 (2021).

[20] Ser-Giacomi, E. et al. Ubiquitous abundance distribution of non-dominant plankton across the global ocean. Nature Ecology & Evolution 2, 1243–1249 (2018).

[21] Chust, G., Irigoien, X., Chave, J. & Harris, R. P. Latitudinal phytoplankton distribution and the neutral theory of biodiversity. Global Ecology and Biogeography 22, 531–543 (2013).

[22] Preston, F. W. The commonness, and rarity, of species. Ecology 29, 254–283 (1948).

[23] Magurran, A. E. Measuring Biological Diversity (Blackwell Publishing, Oxford, 2004).

[24] McGill, B. J. et al. Species abundance distributions: moving beyond single prediction theories to integration within an ecological framework. Ecology Letters 10, 995–1015 (2007).

[25] Brown, J. H., Gillooly, J. F., Allen, A. P., Savage, V. M. & West, G. B. Toward a metabolic theory of ecology. Ecology 85, 1771–1789 (2004).

[26] Marquet, P. A. et al. Scaling and power-laws in ecological systems. Journal of Experimental Biology 208, 1749–1769 (2005).

[27] Harte, J. Maximum entropy and ecology: a theory of abundance, distribution, and energetics (Oxford University Press, Oxford, UK, 2011).

[28] Hubbell, S. P. The unified neutral theory of biodiversity and biogeography (Princeton University Press, Princeton, NJ, 2001).

[29] Volkov, I., Banavar, J. R., Hubbell, S. P. & Maritan, A. Neutral theory and relative species abundance in ecology. Nature 424, 1035–1037 (2003).

[30] Alonso, D. & McKane, A. J. Sampling Hubbell’s neutral theory of biodiversity. Ecology Letters 7, 901–910 (2004).

[31] Armstrong, R. A. Grazing limitation and nutrient limitation in marine ecosystems: steady state solutions of an ecosystem model with multiple food chains. Limnology and Oceanography 39, 597–608 (1994).

[32] Follows, M. J., Dutkiewicz, S., Grant, S. & Chisholm, S. W. Emergent biogeography of microbial communities in a model ocean. Science 315, 1843–1846 (2007).

[33] Ward, B. A., Dutkiewicz, S., Jahn, O. & Follows, M. J. A size-structured food-web model for the global ocean. Limnology and Oceanography 57, 1877–1891 (2012).

[34] Sal, S., López-Urrutia, Á., Irigoien, X., Harbour, D. S. & Harris, R. P. Marine microplankton diversity database: Ecological Archives E094-149. Ecology 94, 1658 (2013).

[35] Dutkiewicz, S., Follows, M. J. & Bragg, J. G. Modeling the coupling of ocean ecology and biogeochemistry. Global Biogeochemical Cycles 23, GB4017 (2009).

[36] Dutkiewicz, S., Ward, B., Monteiro, F. & Follows, M. Interconnection of nitrogen fixers and iron in the Pacific Ocean: Theory and numerical simulations. Global Biogeochemical Cycles 26, GB1012 (2012).

[37] Dutkiewicz, S. et al. Capturing optically important constituents and properties in a marine biogeochemical and ecosystem model. Biogeosciences 12, 4447–4481 (2015).

[38] Marshall, J., Adcroft, A., Hill, C., Perelman, L. & Heisey, C. A finite-volume, incompressible Navier Stokes model for studies of the ocean on parallel computers. Journal of Geophysical Research: Oceans 102, 5753–5766 (1997).

[39] Wunsch, C. & Heimbach, P. Practical global oceanic state estimation. Physica D: Nonlinear Phenomena 230, 197–208 (2007).

[40] Kuhn, A. et al. Temporal and spatial scales of correlation in marine phytoplankton communities. Journal of Geophysical Research: Oceans 124, 9417–9438 (2019).

[41] Dutkiewicz, S., Boyd, P. W. & Riebesell, U. Exploring biogeochemical and ecological redundancy in phytoplankton communities in the global ocean. Global Change Biology 27, 1196–1213 (2021).

[42] Clauset, A., Shalizi, C. R. & Newman, M. E. Power-law distributions in empirical data. SIAM Review 51, 661–703 (2009).

[43] Ewens, W. J. The sampling theory of selectively neutral alleles. Theoretical Population Biology 3, 87–112 (1972).

[44] Burnham, K. P. & Anderson, D. R. Multimodel inference: understanding AIC and BIC in model selection. Sociological Methods & Research 33, 261–304 (2004).

[45] McGill, B. J. A test of the unified neutral theory of biodiversity. Nature 422, 881–885 (2003).

[46] Chave, J. Neutral theory and community ecology. Ecology Letters 7, 241–253 (2004).

[47] Alonso, D., Etienne, R. S. & McKane, A. J. The merits of neutral theory. Trends in Ecology & Evolution 21, 451–457 (2006).

[48] Adler, P. B., HilleRisLambers, J. & Levine, J. M. A niche for neutrality. Ecology Letters 10, 95–104 (2007).

[49] Utermöhl, H. Zur Vervollkommung der quantitativen Phytoplankton-Methodik. Internationale Vereinigung für Theoretische und Angewandte Limnologie: Mitteilungen 9, 1–38 (1958).

[50] Polz, M. F. & Cavanaugh, C. M. Bias in template-to-product ratios in multitemplate PCR. Applied and Environmental Microbiology 64, 3724–3730 (1998).

[51] Bradley, I. M., Pinto, A. J. & Guest, J. S. Design and evaluation of Illumina MiSeq-compatible, 18S rRNA gene-specific primers for improved characterization of mixed phototrophic communities. Applied and Environmental Microbiology 82, 5878–5891 (2016).

[52] Gong, W. & Marchetti, A. Estimation of 18S gene copy number in marine eukaryotic plankton using a next-generation sequencing approach. Frontiers in Marine Science 6, 219 (2019).

[53] Menemenlis, D. et al. ECCO2: high resolution global ocean and sea ice data synthesis. Mercator Ocean Quarterly Newsletter 31, 13–21 (2008).

[54] Yodzis, P. & Innes, S. Body size and consumer-resource dynamics. The American Naturalist 139, 1151–1175 (1992).

[55] Martinez, N. D. Allometric trophic networks from individuals to socio-ecosystems: consumer–resource theory of the ecological elephant in the room. Frontiers in Ecology and Evolution 8, 92 (2020).

[56] Menden-Deuer, S. & Lessard, E. J. Carbon to volume relationships for dinoflagellates, diatoms, and other protist plankton. Limnology and Oceanography 45, 569–579 (2000).

[57] Scott, D. W. On optimal and data-based histograms. Biometrika 66, 605–610 (1979).

[58] Freedman, D. & Diaconis, P. On the histogram as a density estimator: L2 theory. Zeitschrift für Wahrscheinlichkeitstheorie und Verwandte Gebiete 57, 453–476 (1981).

[59] Karlin, S. & McGregor, J. Addendum to a paper of W. Ewens. Theoretical Population Biology 3, 113–116 (1972).

[60] Tavaré, S. & Ewens, W. J. in Multivariate Ewens distribution (eds Johnson, N. L., Kotz, S. & Balakrishnan, N.) Discrete Multivariate Distributions 232–246 (Wiley, New York, 1997).

[61] Akaike, H. A new look at the statistical model identification. IEEE Transactions on Automatic Control 19, 716–723 (1974).

[62] Sal, S., López-Urrutia, Á., Irigoien, X., Harbour, D. S. & Harris, R. P. Marine microplankton diversity database. Figshare (2013). URL 10.6084/m9.figshare.c.3306033.

