## Supplementary Information for: Invariant scaling of the Species Abundance Distribution in observed and simulated marine plankton communities for "Invariant scaling of the Species Abundance Distribution in observed and simulated marine plankton communities"

[1]MIT-WHOI Joint Program in Oceanography/Applied Ocean Science and Engineering, Cambridge and Woods Hole, MA, USA.
[2]Biology Department, Woods Hole Oceanographic Institution, Woods Hole, MA, USA.
[3]Department of Earth, Atmospheric, and Planetary Sciences, Massachusetts Institute of Technology, Cambridge, MA, USA.
[4]Instituto de Física Interdisciplinar y Sistemas Complejos (IFISC, CSIC–UIB), Universitat de les Illes Balears, Palma de Mallorca, Islas Baleares, Spain.
[5]Center for Sustainability Science and Strategy, Massachusetts Institute of Technology, Cambridge, MA, USA.

#### Supplementary Information

**Supplementary Table 1** Cruise metadata for the marine microplankton diversity database locations shown in main text Fig. 1, compiled from the database metadata [1]. Sampling period gives the first and last sample dates for each cruise, with dates reported in YYYY-MM-DD. Surface observations are the number of surface sample records in the respective cruise. Total observations are all sample records across depths. Latitude and longitude ranges summarize the sampled extent.

| Cruise | Sampling period | Surface obs. | Total obs. | Lat. (°) | Lon. (°) |
|---|---|---:|---:|---|---|
| AMT1 | 1995-09-25 to 1995-10-24 | 25 | 50 | [-50.8, 48.9] | [-57.4, -9.1] |
| AMT2 | 1996-04-23 to 1996-05-21 | 25 | 53 | [-47.6, 48.7] | [-55.6, -8.1] |
| AMT3 | 1996-09-24 to 1996-10-25 | 25 | 76 | [-51.9, 47.4] | [-57.9, -18.2] |
| AMT4 | 1997-04-21 to 1997-05-23 | 27 | 54 | [-51.0, 48.1] | [-57.3, -14.0] |
| AMT5 | 1997-09-18 to 1997-10-15 | 24 | 48 | [-46.5, 48.0] | [-56.8, -13.2] |
| AMT6 | 1998-05-16 to 1998-06-13 | 11 | 31 | [-32.1, 5.9] | [-16.1, 17.9] |
| Benguela | 2000-10-15 to 2000-10-28 | 53 | 54 | [-25.6, -18.8] | [-14.8, -11.0] |
| Bergen | 2000-06-06 to 2000-06-24 | 1 | 46 | 60.3 | 5.2 |
| FISHES | 2001-05-11 to 2001-05-28 | 24 | 25 | [60.1, 65.4] | [-25.5, -8.4] |
| Irminger | 2002-05-11 to 2002-05-22 | 8 | 8 | [61.2, 65.3] | [-37.5, -25.4] |
| l4 | 1992-10-12 to 2001-10-29 | 1 | 360 | 50.2 | -4.2 |
| LongIsland | 2001-02-05 to 2001-03-26 | 1 | 7 | 41.3 | -72.0 |
| NorthSea | 1994-10-08 to 1995-07-11 | 44 | 44 | [52.9, 55.9] | [-1.9, 1.8] |
| Norwegian | 1997-03-22 to 1997-06-05 | 1 | 19 | 66.0 | 2.0 |
| Oregon | 2001-05-18 to 2001-09-10 | 1 | 7 | 44.7 | -124.2 |
| Prime | 1996-06-18 to 1996-06-26 | 9 | 50 | [59.0, 59.2] | [-20.6, -20.0] |
| Sonne | 1997-05-19 to 1997-07-07 | 16 | 111 | [16.0, 18.4] | [56.9, 62.0] |

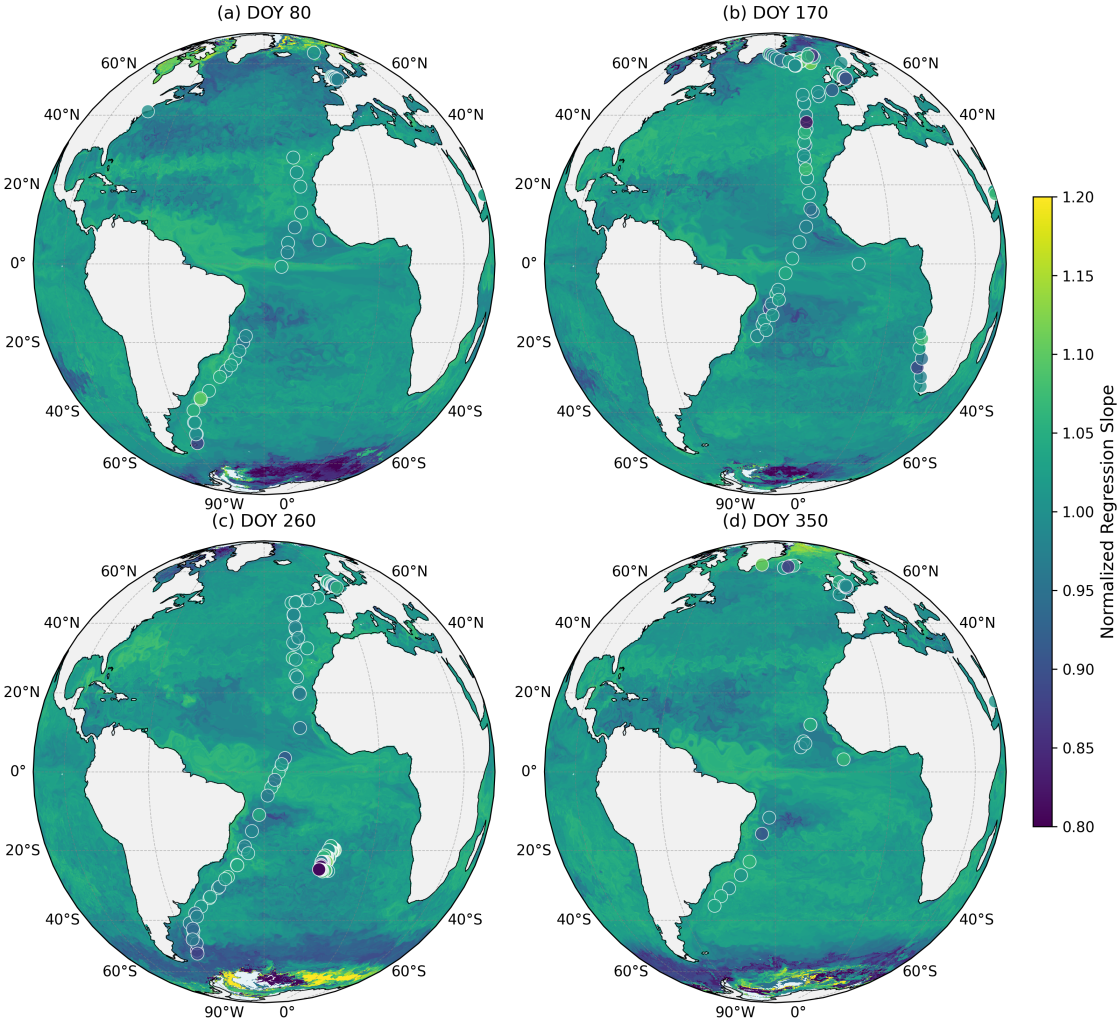


**Supplementary Fig. 1** Global maps of normalized regression slopes of SADs. Slopes describe the relationship between log abundance and log frequency, normalized by the dataset or cruise mean. Observational data are overlaid as scatter points, matched to the closest pixel and day-of-year.

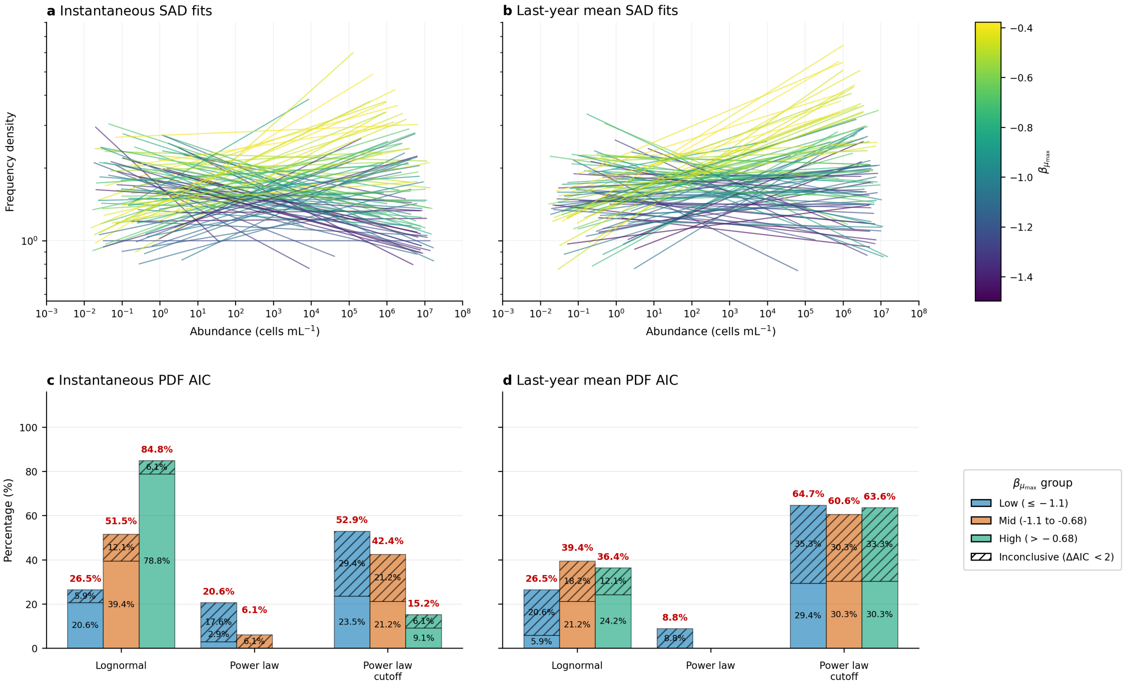


**Supplementary Fig. 2** SAD regressions and best fitting PDF selection for the Armstrong (1994) simulations [2]. To compare with the results in Fig. 4a–e, we plotted both snapshots at the last saved time step and annual averages over the final year. Panels show (a) SAD regression fits for final-time snapshots, (b) SAD regression fits for final-year mean abundances, (c) best fitting PDF selection for final-time snapshots, and (d) PDF selection for final-year mean abundances. The community comprises 50 phytoplankton and 50 zooplankton size classes. Phytoplankton diameters span 0.5–130 $\mu$m (geometric progression with ratio 1.12), and zooplankton diameters are taken as 10× the corresponding phytoplankton sizes (5–1300 $\mu$m). Dynamics include Monod phytoplankton growth and piecewise-linear grazing; total nutrient is fixed at $T = 10$ mmol N m$^{-3}$ and food chains are independent. Base values for the smallest class follow the Armstrong standard case: $\mu_{\max,0} = 1.4$ d$^{-1}$, $H_{\max,0} = 1.4$ d$^{-1}$, $\delta_0 = 0.168$ d$^{-1}$, with allometric exponents $\beta_{H_{\max}} = \beta_\delta = -0.75$ (other parameters are size-independent: $K_s = 0.1$, $X = 0.016$, $K_p = 2.0$, $y = 0.4$). Biomass is converted to abundance (cells mL$^{-1}$) using carbon–volume scaling [3, 4], with values $< 0.005$ cells mL$^{-1}$ set to zero. For each random draw of $\beta_{\mu_{\max}}$, pooled P+Z abundances are binned on a log scale (10 bins), and a straight line is fit in log–log space to the nonzero bins; line color indicates $\beta_{\mu_{\max}}$. For PDF selection, pooled P+Z abundances are fit to lognormal, power-law, and power-law with exponential cutoff distributions, and AIC values are compared. Bars show the percentage of simulations in which each PDF is best supported within low, mid, and high $\beta_{\mu_{\max}}$ groups; hatched segments indicate inconclusive cases with $\Delta$AIC $< 2$, and red labels show the total percentage including conclusive and inconclusive cases.

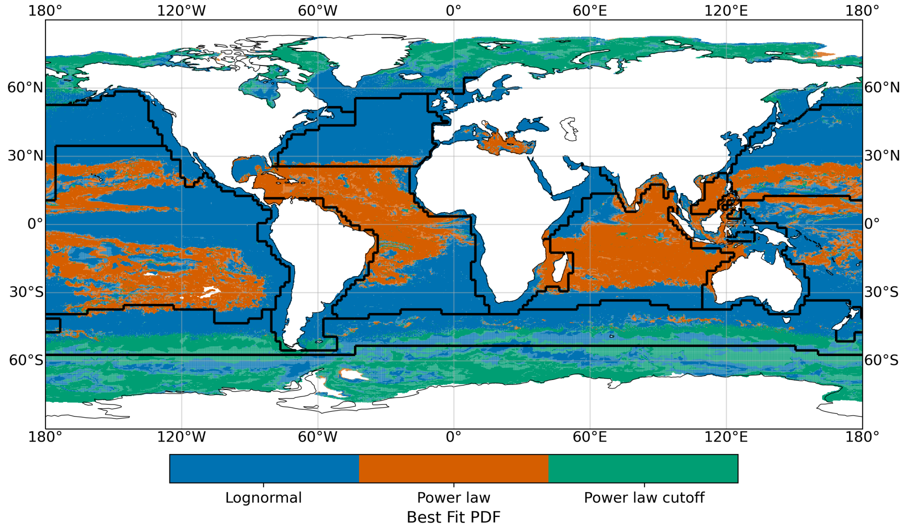


**Supplementary Fig. 3** Global maps of best fitting PDFs determined by AIC across grid cells in Darwin. Inconclusive fits ($\Delta$AIC $< 2$) are shown with reduced opacity. Black contours indicate biome boundaries.

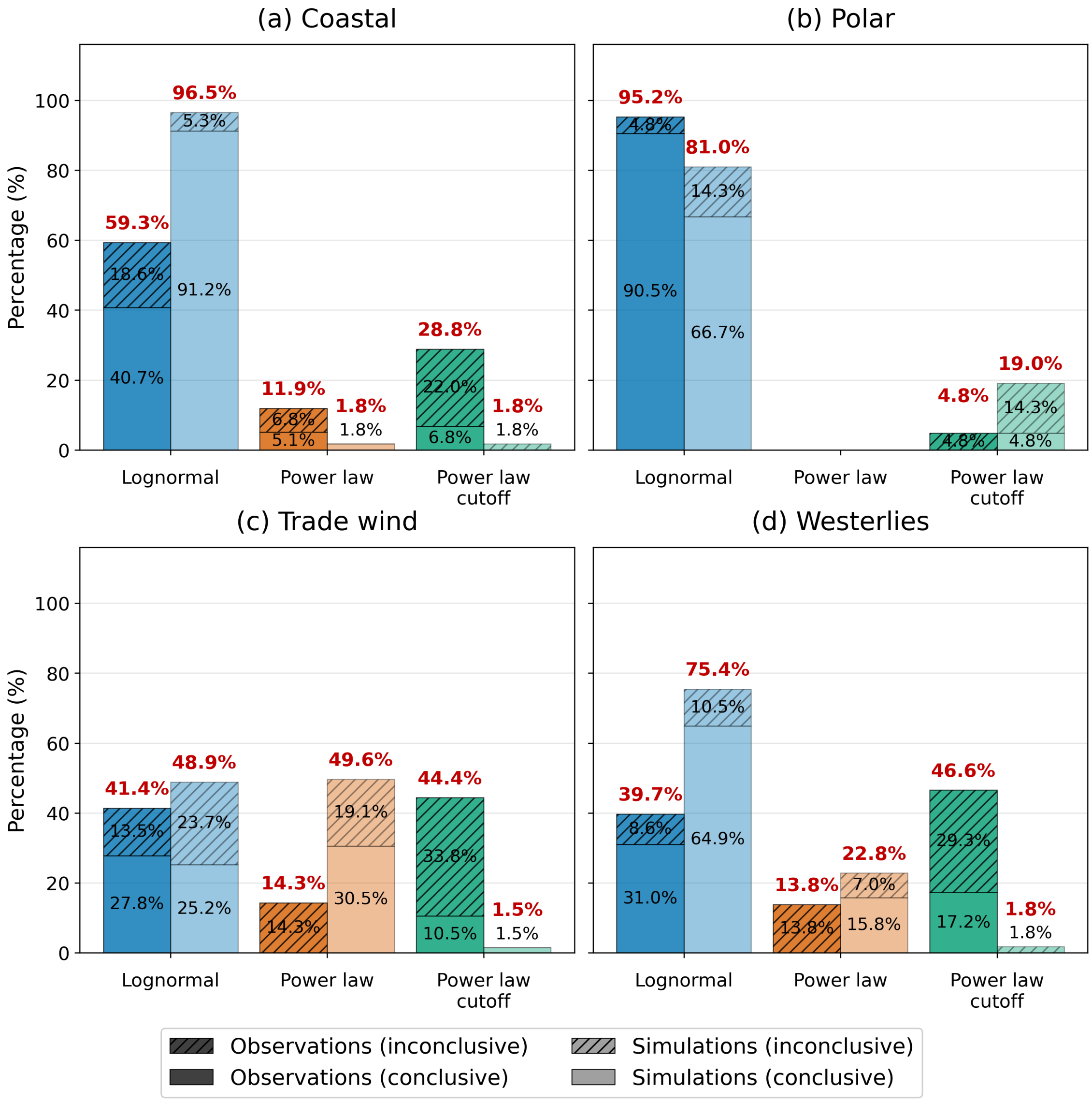


**Supplementary Fig. 4** Percentage of observational and station-matched simulated samples best fit by each probability density function (PDF) type across four marine biomes. (a–d) Coastal, Polar, Trade wind, and Westerlies biomes, respectively. Bars show the percentage of successful fits assigned to each PDF type: lognormal, power law, and power law with cutoff. Observational samples are shown with darker bars and station-matched simulated samples with lighter bars. Conclusive fits ($\Delta$AIC $\geq 2$) are shown as solid bar segments, while inconclusive fits ($\Delta$AIC $< 2$) are hatched and stacked above the corresponding conclusive segments. Black percentage labels indicate the individual conclusive and inconclusive contributions, and bold red labels above each bar indicate the total percentage for that PDF type.

### References

[1] Sal, S., López-Urrutia, Á., Irigoien, X., Harbour, D. S. & Harris, R. P. Marine microplankton diversity database: Ecological Archives E094-149. Ecology **94**, 1658 (2013).

[2] Armstrong, R. A. Grazing limitation and nutrient limitation in marine ecosystems: steady state solutions of an ecosystem model with multiple food chains. Limnology and Oceanography **39**, 597–608 (1994).

[3] Menden-Deuer, S. & Lessard, E. J. Carbon to volume relationships for dinoflagellates, diatoms, and other protist plankton. Limnology and Oceanography **45**, 569–579 (2000).

[4] Ward, B. A., Dutkiewicz, S., Jahn, O. & Follows, M. J. A size-structured food-web model for the global ocean. Limnology and Oceanography **57**, 1877–1891 (2012).
